# A High Throughput Assay for Measuring Secreted Protein Based on a *de novo* Fluorescent Reporter Reveals Regulatory and Structural Insights in *Salmonella* Type Three Secretion System

**DOI:** 10.1101/2025.01.17.633628

**Authors:** Samuel Alexander Leach, Jordan Scott Summers, Edward Wen, Danielle Tullman-Ercek

## Abstract

Intracellular protein production in bacteria is limited by the need for lysis and costly purification. A promising alternative is to engineer the host organism for protein secretion. While the *Salmonella enterica* serovar Typhimurium (*S*. Typhimurium) Type 3 Secretion System (T3SS) has been utilized for protein secretion, its study and eventual applicability for recombinant protein production is constrained by the lack of high-throughput assays to quantitatively measure secretion titer. Developing such assays is challenging, as proteins must remain unfolded for secretion, limiting the use of several common reporter proteins. In this work, we develop a high-throughput secretion assay using mini-Fluorescent Activating Protein (mFAP). mFAP forms a chromophore only upon addition of an exogenous substrate, allowing secretion and subsequent fluorescence detection. We demonstrate mFAP secretion via the T3SS with an N-terminal secretion tag and show that the fluorescent signal in the secreted fraction is rapid and linear over three orders of magnitude. Using this assay, we screen *S*. Typhimurium strains with secretion-enhancing mutations, identifying a constitutively active strain and reveal temporally controlled secretion dynamics. We also show that this assay may be applicable to other secretion systems, providing a universal tool for tracking heterologous protein secretion.

## Introduction

Heterologous proteins are utilized in products which generate revenues exceeding $100 billion (“Global Bioengineered Protein Drugs Market Growth Analysis Report,” n.d.). Bacterial hosts such as *Escherichia coli* offer advantages for protein production, including genetically tractable systems, high titers, fast growth, and high cell density in commercial media (Rosano & Ceccarelli, 2014). However, these hosts typically retain proteins intracellularly, complicating purification by requiring cell lysis, and high intracellular expression also increases toxicity risks to the cells which can limit titer (Lilie, Schwarz, & Rudolph, 1998). A promising strategy is to engineer bacterial hosts to secrete proteins, thus simplifying purification and reducing toxicity. Bacteria possess various secretion systems, many of which are well studied and have been adapted for heterologous protein secretion, including the Type III Secretion System (T3SS) (Burdette, Leach, Wong, & Tullman-Ercek, 2018). While several bacterial secretion systems have been adapted for this purpose (Kotzsch et al., 2011), their mechanisms remain poorly understood and increasing secretion titer to rival conventional protein production strategies remains a major challenge.

Advancing our understanding of secretion mechanisms and increasing yield requires high-throughput technologies for screening secretion titers. For example, Lesser et al. used a high-throughput screen to uncover that *Shigella flexneri* effectors were efficiently secreted via a T3SS without their associated chaperones, revealing a chaperone-independent secretion mechanism (Ernst, Reeves, Ramseyer, & Lesser, 2018). High-throughput screening is also essential for leveraging large mutant libraries, such as transposon mutagenesis libraries and the Keio knockout collection, to enhance secretion. In several studies, researchers used these tools to improve heterologous protein secretion in *E. coli* lab strains (Natarajan, Haitjema, Lee, Boock, & DeLisa, 2017). To continue these successes, there is a need for high throughput assays for accurate quantification of secreted protein.

Several strategies are used to detect and quantify secreted heterologous proteins, including enzymatic reporter assays. These assays rely on secreted enzymes that catalyze reactions to produce measurable signals, such as amino-peptidase and β-1,4-mannanase in Gram-positive bacteria (Guan et al., 2016; Lin et al., 2015). However, these assays can be challenging to apply across contexts. Some enzymes, such as alkaline phosphatase and beta-lactamase, require cofactors or periplasmic folding, which may not occur if secretion bypasses the periplasm (Elfageih et al., 2020; Minsky, Summers, & Knowles, 1986). Enzyme activity is also influenced by pH, salt concentration, and environmental factors (Metcalf et al., 2016; Srinivasan, Kantae, & Robinson, 2020), further complicating the correlation between activity and protein concentration under varying conditions.

Fluorescent reporter assays overcome many challenges of enzymatic assays, offering rapid signal maturation, accurate measurement across conditions, and high-throughput compatibility with plate readers. However, common fluorescent proteins like GFP are rarely used in secretion assays since they cannot unfold and refold during secretion (Akeda & Galán, 2005; Jaumouillé, Francetic, Sansonetti, & Tran Van Nhieu, 2008; Radics, Königsmaier, & Marlovits, 2013). To address this, secretable reporters that fluoresce with exogenous substrates—such as TEM-1 β-lactamase, FlAsH-tetracysteine, and split GFP assays—have been developed (Enninga, Mounier, Sansonetti, & Nhieu, 2005; Haitjema et al., 2014; Mills, Baruch, Aviv, Nitzan, & Rosenshine, 2013; Van Engelenburg & Palmer, 2010; Zhu et al., 2011). While useful for identifying secretion substrates and tracking secretion dynamics, these assays require significant optimization, are typically performed in minimal media to avoid high background, and can be costly due to substrate expense, limiting their use in large-scale studies.

Recently, researchers in the Baker lab designed, built, and tested a *de novo* protein consisting of a single beta barrel which is capable of selectively binding the inexpensive substrate DFHBI to produce a fluorescent signal, called the mini Fluorescence Activating Protein (mFAP) (Dou et al., 2018). This protein can be visualized *in vitro* and *in vivo* after the addition of the substrate. Because this protein can produce a fluorescent signal only through the addition of a substrate, we hypothesized that the mFAP protein may also be suitable as a reporter in a fluorescence-based secretion assay.

The T3SS of *Salmonella enterica* serovar Typhimurium (*S*. Typhimurium) is ideal for engineering high-titer heterologous protein secretion. This multiprotein structure spans both membranes and secretes pathogenic effectors directly into host cells, bypassing the periplasm. With N-terminal secretion tags, heterologous proteins can be secreted extracellularly into the media under standard culture conditions. The T3SS’s well-characterized structure, regulation, and non-essential nature make it a prime target for engineering without affecting growth. While progress has been made in increasing secretion titers (Glasgow, Wong, & Tullman-Ercek, 2017; Kevin J Metcalf, Finnerty, Azam, Valdivia, & Tullman-Ercek, 2014), challenges remain, requiring high-throughput strategies to engineer structural proteins (Burdette et al., 2024), rewire regulation (Song et al., 2017a), and optimize secretion tags.

In this study, we develop mFAP into a fluorescence assay for measuring secretion titer. We show that mFAP can be secreted through the T3SS of *S*. typhimurium with a T3SS N-terminal secretion tag and fluoresces extracellularly upon DFHBI addition. We establish its dynamic detection range, demonstrate its utility by screening combinatorial genomic mutants, and explore secretion dynamics. Finally, we show that mFAP can detect secretion through other mechanisms, highlighting its potential as a generalizable assay for engineering secretion systems and enabling large-scale studies.

## Results

### mFAP can be used to assay secretion through the T3SS

We used the well-characterized T3SS of *S.* Typhimurium to identify a reporter that can be secreted while producing a stable fluorescent signal. The recently developed *de novo* protein, mFAP, consists of a single β-barrel which specifically binds to the substrate DFHBI to produce a fluorescent signal, making it an ideal candidate for secretion studies. We hypothesized that mFAP could be secreted through the T3SS and generate fluorescence upon DFHBI addition.

To test if secreted mFAP fluoresces extracellularly, we incorporated mFAP version 2a (Dou et al., 2018) into our platform for secreting heterologous proteins via the T3SS of *S.* Typhimurium (Kevin J Metcalf et al., 2014). In this platform, mFAP2a is fused to the C-terminus of SptP^167^, an N-terminal secretion tag for T3SS secretion, with 2xFLAG and 6xHis tags for purification and detection. The fusion (hereafter referred to as SptP-mFAP) is produced via expression from a “secretion vector” under the native T3SS promoter P*_sic_*. An “activation vector” provides IPTG-inducible *hilA* expression via the P*_lac_* promoter, and HilA in turn activates T3SS assembly and secretion. We transformed *S*. Typhimurium strain ASTE13 with the secretion and activation vectors, used IPTG to induce T3SS assembly followed by SptP-mFAP secretion, and collected secreted samples after 8 hours. The samples were incubated with 100 µM DFHBI, and fluorescence was measured (Fig. 1A).

**Figure 1:**
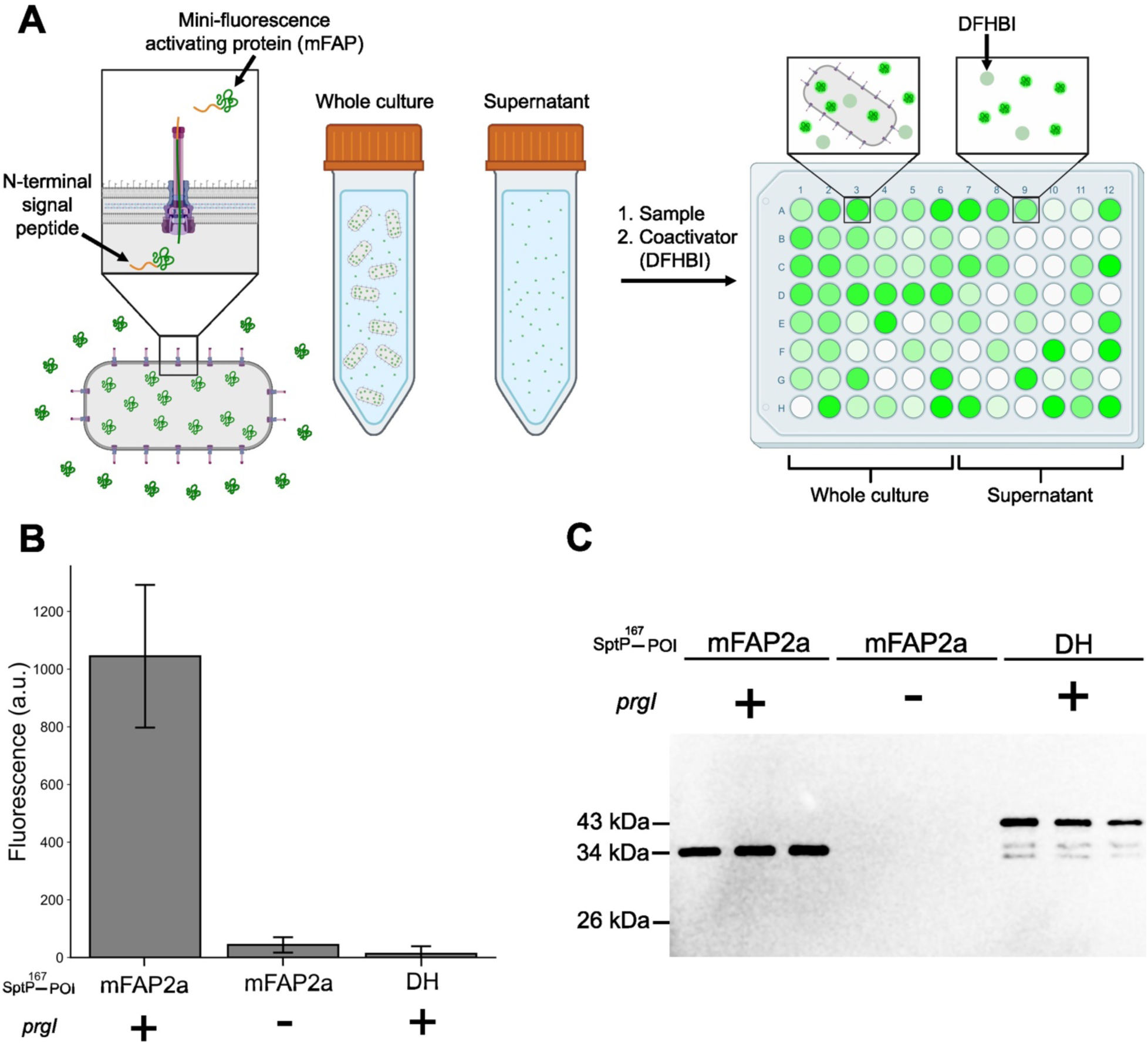
SptP-mFAP2a produces a fluorescent signal after secretion via the T3SS. A) *S*. Typhimurium strains secreting mFAP are separated into whole culture and supernatant by centrifugation. Each sample is added to a 96-well plate, incubated with 100 μM DFHBI and fluorescence is read on a plate reader. B) A strain harboring the SptP-mFAP2a secretion plasmid was grown for 8 hours, a sample of the secreted fraction was collected after centrifugation, and mFAP fluorescence was induced with DFHBI. The fluorescence of the induced samples was measured alongside secretion samples from a secretion incompetent strain (Δ*prgI*), and a strain harboring the non-fluorescent SptP-DH secretion plasmid. Error bars reflect the standard error the mean of three biological replicates. C) Western blot of the secreted fraction from 1B. Figure 1A created in BioRender. Summers, J. (2025) https://BioRender.com/c83r875

To ensure the fluorescence was from mFAP and not another protein in the secreted fraction, we used a construct replacing the gene encoding mFAP with the gene encoding a non-fluorescent DH domain from human intersectin (Ahmad & Lim, 2010). Despite detecting SptP^167^-DH in the extracellular space, fluorescence remained near background (Fig. 1B). To confirm that SptP-mFAP secretion was T3SS-dependent, we measured fluorescence and probed for FLAG tags in a strain lacking PrgI, a key T3SS component (Kubori, Sukhan, Aizawa, & Galán, 2000). This secretion-incompetent strain showed minimal fluorescence (Fig. 1B) and no detectable protein was observed on the western blot (Fig. 1C), confirming that SptP-mFAP secretion required the T3SS. These results confirm that SptP-mFAP is specifically secreted via the T3SS and produces a fluorescent signal.

### The mFAP assay is rapid, stable, and has a wide detection range

Having established that fluorescence in the media is a result of T3SS secretion of mFAP, we aimed to develop a robust secretion assay usable across multiple contexts. While an mFAP-based assay can be simple and efficient, requiring only a plate reader measurement, we sought to ensure accurate measurements across various secretion titers.

An ideal secretion assay should produce a linear response across a broad range of titers, from <1 mg/L to hundreds of mg/L (Ling, Rahmat, Murad, Mahadi, & Illias, 2017; Natarajan et al., 2017), with a target range of 1–100 mg/L. Therefore, we set out to determine the range of mFAP concentrations across which the fluorescent signal is linear. To determine mFAP’s linear range, we purified SptP-mFAP, assessed its concentration, diluted it into secreted fractions of *S*. Typhimurium cultures without secretion plasmids, and generated fluorescence calibration curves in two types of media. This allows us to mimic the fluorescence of mFAP at exact amounts as if it were secreted along with the rest of the *S*. Typhimurium secretome. We tested No Carbon Essential (NCE) medium with 0.4% glycerol as a model for low background fluorescence and LB-L as a model for higher autofluorescence due to yeast extract components (Maslanka, Kwolek-Mirek, & Zadrag-Tecza, 2018). The mFAP assay showed linear fluorescence from 1 mg/L to 100 mg/L in both media (Fig. 2A), demonstrating compatibility across conditions.

**Figure 2:**
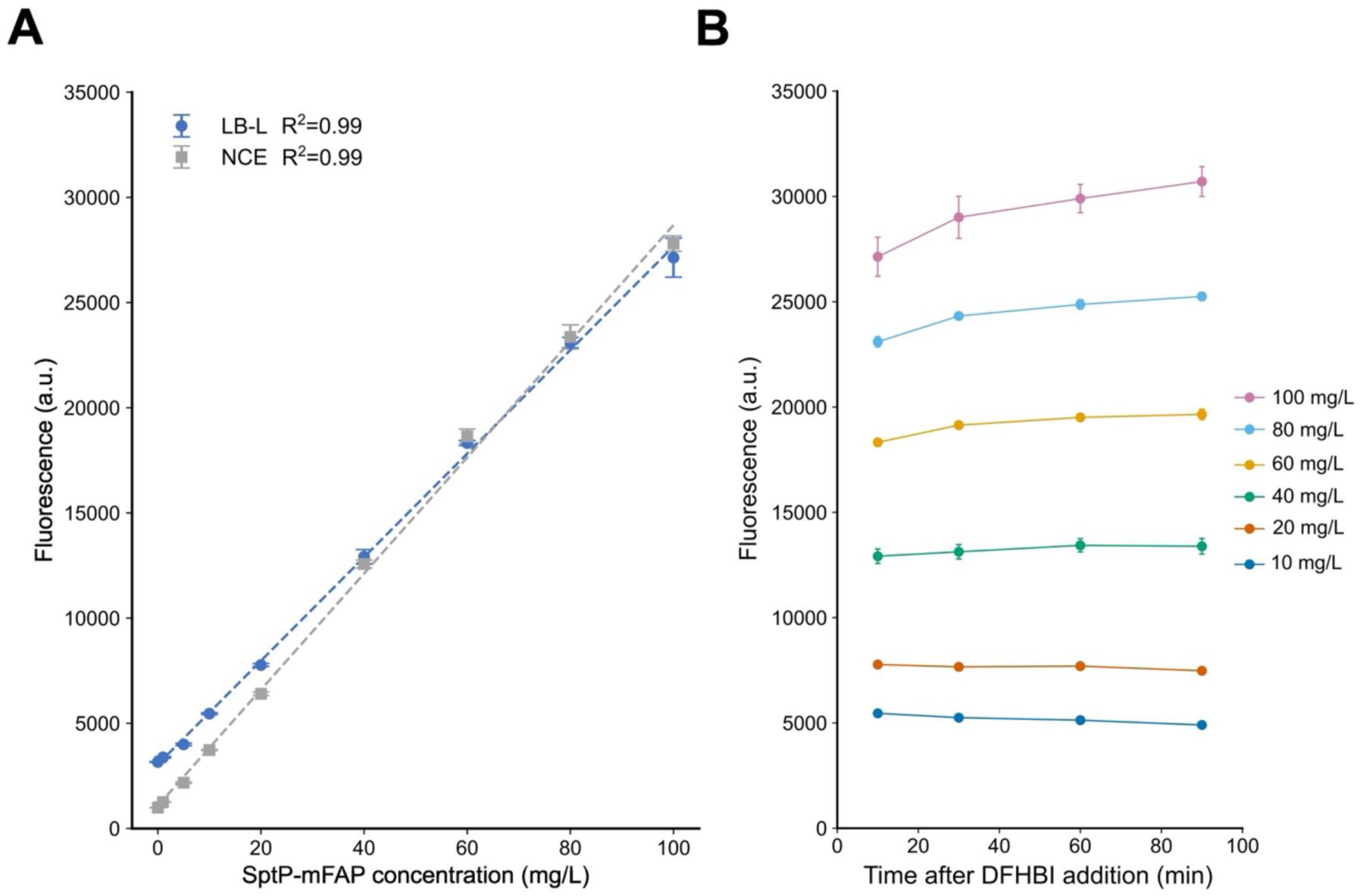
The mFAP assay produces a linear signal and develops rapidly. A) Purified SptP-mFAP (1–100 mg/L) was diluted into the secreted fraction of *S.* Typhimurium strains lacking secretion plasmids. Fluorescence was measured after DFHBI addition, with linear fits generated in rich (LB-L) and defined (NCE) media. R² values indicate fit quality, with error bars showing standard deviation from three technical replicates. B) LB-L samples were measured over 10–90 minutes. Each color represents a different SptP-mFAP concentration (10-100 mg/L), with error bars indicating standard deviation from three technical replicates.

We previously reported that adding potassium phosphate and glycerol to LB-L (enhanced LB-L, hereafter referred to as LB-ES) boosts secretion titer by as much as fivefold (Burdette, Wong, & Tullman-Ercek, 2021a). We set out to validate these findings using the mFAP assay and observed a similar four-to five-fold titer increase in SptP-mFAP strains grown in LB-ES compared to LB-L, consistent with results from semi-quantitative western blot (Supplementary Fig. 1A). Further, a calibration curve with varying concentrations of SptP-mFAP in LB-L or LB-ES showed no substantial change in fluorescence (Supplementary Fig. 1B).

We next set out to determine how quickly the mFAP-DFHBI fluorescent signal develops. Since the complex is more photostable than GFP, which requires chromophore maturation and undergoes photobleaching (Klima et al., 2021), we hypothesized rapid signal maturation. Varying concentrations of SptP-mFAP were added to the secreted fraction of *S*. Typhimurium and fluorescence was induced by addition of 100 μM DFHBI and measured at different time points (Fig. 2B). We observed low percent variation in fluorescence over time for most protein concentrations tested (Supplementary Fig. 1C). While we validated this assay for high titers, it is desirable to measure low quantities, such as native secretion in the µg/L range, as well. The Baker lab recently developed mFAP10, a brighter version of mFAP2a (Klima et al., 2021). When fused to SptP^167^, mFAP10 is at least five times brighter than mFAP2a (Supplementary Fig. 2A). These findings were further validated by measuring fluorescence between mFAP2a and mFAP10 at >1 mg/L. We find that mFAP10 displays more linearity compared to mFAP2a at the same protein concentration, enabling more robust protein detection using mFAP10 at relatively low physiological concentrations (Supplementary Fig. 2B). The quick, stable signal offers flexibility in sample measurement timing, mFAP variant choice, and detection of proteins with lower concentrations in the secreted fraction.

### The mFAP assay is amenable to studying engineering of the T3SS

We further tested mFAP’s reliability as a reporter by fusing it to seven N-terminal secretion tags previously evaluated for heterologous protein secretion by Widmaier et al (Widmaier et al., 2009). In that work, western blotting was used to assess secretion of various biopolymer proteins fused to each of these tags, and they observed that fusions to SptP^167^ generally yielded the highest titers. Here, we assessed the secretion titer conferred by each tag when fused to mFAP, and confirmed that SptP^167^ conferred the highest relative titer, as expected, while the others resulted in less than 20% of its titer (Fig. 3A). Intriguingly, we observed that any tag that enabled secretion of over 10% of the titer achieved with the SptP^167^ fusion also gave consistent results between the mFAP assay and semi-quantitative western blot, while those with titers below 10% of that level were detectable via fluorescence but had no detectable signal on the blot, likely due to being outside the dynamic range for western blotting. The wide dynamic range of the mFAP assay highlights the ability to compare samples across a broad titer range without concentration adjustments using the mFAP assay as compared to traditional, lower-throughput methods such as western blotting.

**Figure 3:**
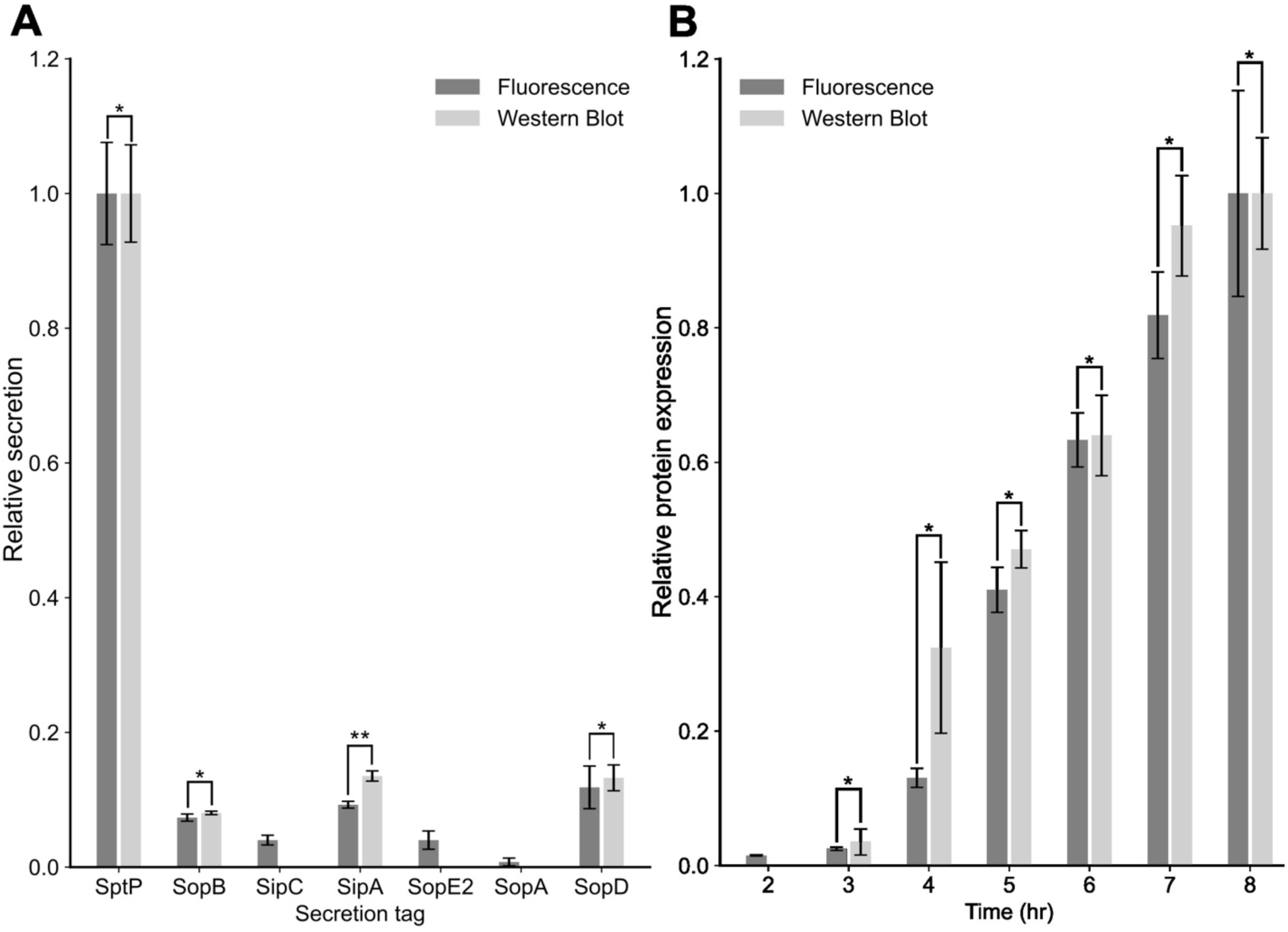
The mFAP assay robustly measures relative protein expression and secretion across different conditions. A) Strains expressing different secretion tags fused to mFAP were compared to SptP-mFAP using fluorescence (dark gray) and semi-quantitative western blot (light gray). Error bars represent standard error from three biological replicates. Significance is marked by * (p < 0.05) and ** (p < 0.01) using Welch’s t-test. B) A strain expressing SptP-mFAP was grown, with whole-culture samples collected over 2–8 hours. Fluorescence and relative protein expression (via western blot) were compared to the 8-hour time point. Error bars represent standard error from three biological replicates, with significance marked by * (p < 0.05).

While we employed mFAP to measure T3SS secretion titer, it may also be possible to assess intracellular protein expression, enabling simultaneous measurement of expression and secretion to calculate secretion efficiency. Understanding secretion efficiency is crucial for optimizing titer and understanding engineering efforts. Since DFHBI crosses membranes (Dou et al., 2018), mFAP should be detectable intracellularly. As a proof of concept, we collected whole culture samples from *S*. Typhimurium expressing SptP-mFAP2a each hour from two to eight hours and separately measured whole culture fluorescence and expressed protein via semi-quantitative western blotting. As expected, we observed an increase in relative protein expression over time for both the mFAP assay and western blotting (Fig. 3B). In addition to measuring secretion efficiency, utilizing mFAP to measure expression is a promising alternative or complement to other assays for protein expression, since the mFAP assay’s signal can be turned on manually and does not require cell lysis as with some enzymatic assays.

### mFAP assay enables identification of optimal genetic background for maximum secretion titer

The ease and reliability of the mFAP assay allows high-throughput secretion assays to test many conditions or strains in a single experiment. We set out to demonstrate this by screening a library of *Salmonella* strains with a collection of genetic, extracellular, and structural enhancements to our T3SS platform.

We and others have developed a variety of plasmid and genomic modifications which enhance secretion titer (Burdette et al., 2024, 2018; Glasgow et al., 2017). Our activation plasmid-based overexpression of the transcription factor HilA directly activates the genes involved with T3SS assembly and secretion (Liang, Burdette, Wong, & Tullman-Ercek, 2022; Kevin J Metcalf et al., 2014). This activation plasmid leads to enhanced secretion titers relative to strains without a plasmid-expressed transcription factor. Recently, we demonstrated that strains harboring needle protein PrgI with the S49R mutation confer higher secretion titers than strains harboring the wild-type PrgI (Burdette et al., 2024). In another study, we demonstrated that removing the needle tip protein SipD also led to higher titers than the wild-type strain with an intact SipD (Glasgow et al., 2017). Additionally, overexpression of the structural regulator InvE could lead to higher secretion titers (Kubori & Galán, 2002), though that has not been directly tested in our platform. Last, the deletion of the gene encoding the protein HilE confers an increased secretion titer (Liang et al., 2022). HilE is a repressor of HilD, which is the known as the “master regulator” of the T3SS since multiple regulatory signals are fed through the activation of HilD to activate the rest of the system (Lou, Zhang, Piao, & Wang, 2019).

While single modifications have been assessed, it is not known whether these enhancements are additive and whether they can be combined to further improve titer. By employing the mFAP assay, studying combinations of these enhancements is feasible in a high throughput fashion. We therefore constructed strains with different combinations of these enhancements – *prgI::S49R*, 11*sipBCD* (Park et al., 2018) (which removes the entire tip complex rather than just SipD), 11*invE*, and 11*hilE*. We then added the SptP-mFAP secretion plasmid and either no activation plasmid or the *hilA* activation plasmid. Finally, we grew these strains in either LB-L or LB-ES and measured protein expression and secretion titer.

These trends align with previous findings (Burdette et al., 2024, 2021a; Glasgow et al., 2017; Liang et al., 2022; Kevin J Metcalf et al., 2014), confirming the assay’s reliability. First, as expected, the *hilA* activation plasmid in general leads to higher titers as compared to strains without an activation plasmid (Fig. 4A). Second, the removal of the tip complex Δ*sipBCD* leads to higher secretion titers than the other structural mutations, *prgI::S49R* and Δ*invE* (Fig. 4B). This agrees with previous findings that showed the removal of SipD led to a greater increase in secretion than the *prgI::S49R* mutation individually (Glasgow et al., 2017). Third, growing in LB-ES mostly uniformly increased titer in all strains (Fig. 4). This matches previous results showing that the enhanced formulation was additive within a strain deleted for Δ*sipD* (Burdette et al., 2021).

**Figure 4:**
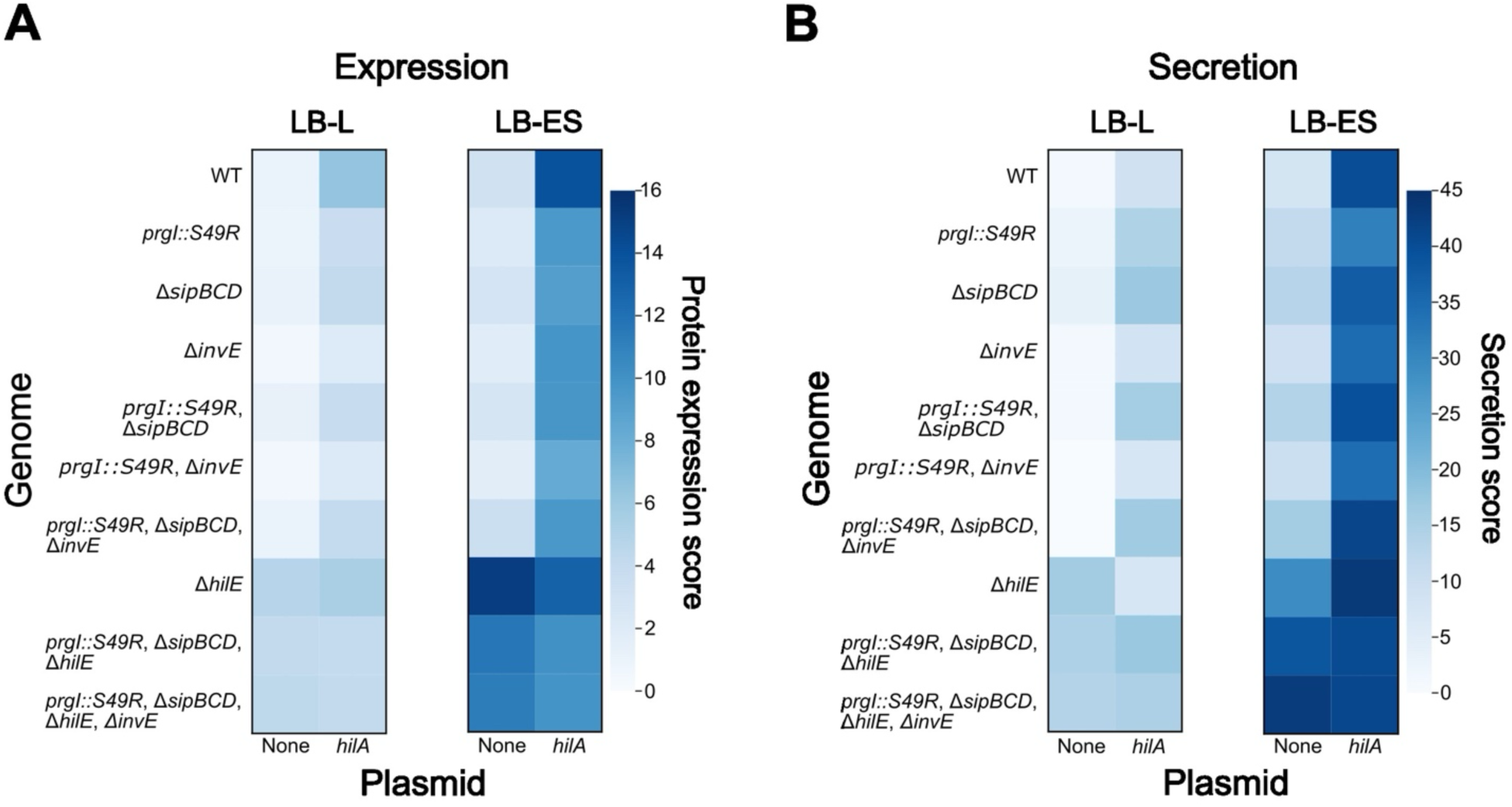
The mFAP assay identifies a constitutively active secretion strain. Strains harboring combinations of genomic mutations (*prgI::S49R*, Δ*sipBCD*, Δ*invE*, and Δ*hilE*, left axis) were transformed with the SptP-mFAP secretion plasmid and either no additional plasmid or the activation plasmid (bottom axis). Strains were grown in either base LB-L or LB-ES, and expression and secretion samples were collected and measured with the mFAP assay. Relative expression and secretion scores are relative to the WT, no plasmid, base LB-L condition and reflect the average of three biological replicates. A) Protein expression data. B) Protein secretion data.

The combination of all structural modifications and Δ*hilE* led to secretion titers comparable to those conferred by the *hilA* activation plasmid (Fig. 4B). The highest titer—about 40-fold higher than the baseline—was achieved in a strain with all genomic modifications but no activation plasmid, grown in LB-ES. Notably, this strain reached ∼80 mg/L (as compared to a standard curve (Fig. 2A)) without synthetic induction. This strain allows for production at high titer without an inducer, which is an attractive feature for commercial protein production.

The mFAP assay can be used for measurements of both protein expression and secretion, which when combined enables determination of secretion efficiency. If secretion titer rises in step with expressed protein, efficiency remains unchanged. If secretion increases without higher expression, efficiency has improved. To maximize titer, secretion efficiency must be optimized. Examination of our data revealed that structural changes to the T3SS secretion apparatus (Δ*sipBCD*, *prgI::S49R*, Δ*invE*) confer reduced or equal mFAP protein expression levels but increase secretion titer compared to WT, indicating improved efficiency (Fig. 4). Regulatory changes (+*hilA*, Δ*hilE*) raise both protein expression and secretion, though secretion increases more, showing enhanced protein expression and efficiency (Fig. 4A).

### The mFAP assay enables a detailed study of secretion dynamics

The mFAP assay’s high throughput ability also enables tracking secretion dynamics over time. While transcriptional data suggest T3SS activation varies with regulatory changes e.g. *hilA* overexpression (Metcalf et al., 2014), we do not have direct evidence showing how these regulatory changes affect secretion over time. The mFAP assay simplifies analyzing multiple time points across conditions, allowing us to answer these questions for the first time.

Using the mFAP assay, we examined how T3SS regulation affects secretion over time. Strains with synthetic *hilA* expression or Δ*hilE* harboring the SptP-mFAP secretion plasmid were grown, and samples were collected hourly between 4 and 10 hours (Fig. 5A). Secretion increased sharply after 6 hours with *hilA* expression, while the Δ*hilE* strain showed delayed but rapid secretion, consistent with derepression of native HilA. These time series trends, enabled by the throughput of the mFAP assay, show that the secretion activity tracks the observed transcriptional activity of the T3SS (Liang et al., 2022). Notably, while both Δ*hilE* and *hilA* overexpression lead to lower cell density compared to WT over the time course (Fig. 5B), we nonetheless observe higher relative secretion titer from these strains. These results are encouraging because this assay permits us to relate transcriptional activity to secretion activity for the first time.

**Figure 5:**
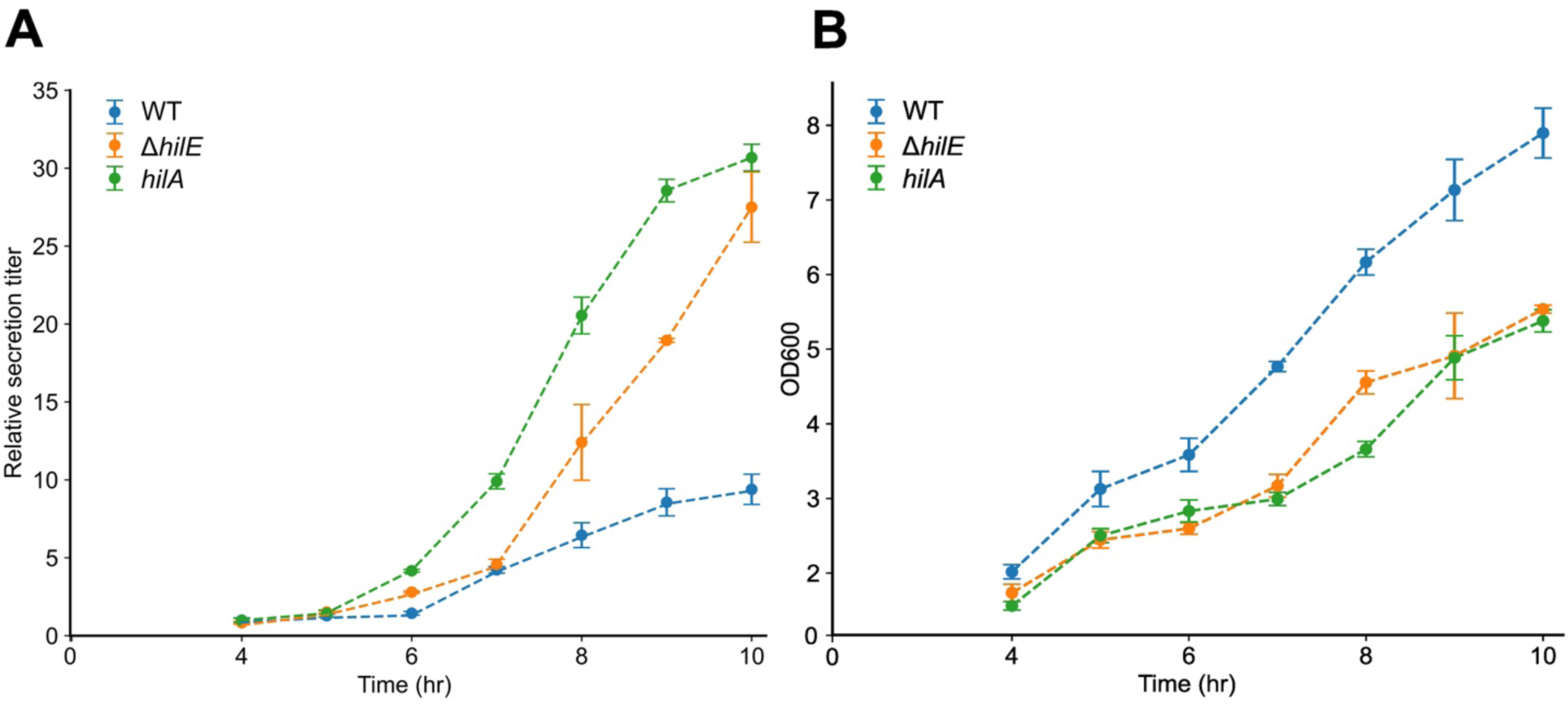
Measuring secretion dynamics over time. A) Strains harboring the SptP-mFAP secretion plasmid and either no additional plasmid, the *hilA* expression plasmid, Δ*hilE* or were grown, and secretion samples were taken every hour between 4 and 10 hours. Fluorescence of the secretion samples were measured with the mFAP assay. Error bars represent the standard error of the mean for three biological replicates. B) The density of the cultures as measured by OD_600_

### The mFAP assay can be used to measure secretion through other secretory mechanisms

While we demonstrated mFAP as a reporter for the T3SS, many bacterial systems are engineered to secrete heterologous proteins (Burdette et al., 2018). Using mFAP as a general reporter across these systems would be advantageous for engineering efforts. Given its small size and single beta-barrel structure, we hypothesized the mFAP assay could be adapted to other secretion pathways and remain detectable through extracellular fluorescence.

We tested this hypothesis with YebF and OsmY from *E. coli*. Both of these are proteins secreted extracellularly through unclear mechanisms, which are used as tags to direct heterologous proteins to the extracellular space (Kotzsch et al., 2011; Zhang, Brokx, & Weiner, 2006). We designed a genetic fusion encoding YebF and OsmY at the N-terminus of mFAP and expressed the genes of a plasmid in *E. coli* BL21(DE3) under the T7 promoter. After incubating the secreted fraction with DFHBI, we detected fluorescence (Fig. 6) and confirmed the presence of the proteins via western blot (Supplementary Fig. 3). We confirmed the fluorescence detected was due to selective secretion by blotting against mFAP fused to the *S*. Typhimurium SptP^167^ secretion tag, which does not confer secretion in *E. coli* (Supplementary Figure 3). The lack of SptP-mFAP in the secreted fraction supports the observed YebF/OsmY-mFAP fluorescence is due to the presence of these unique secretion-inducing fusions. Our success in detecting mFAP secreted with multiple fusion partners and fluorescence indicates that mFAP can most likely be a generalizable reporter capable of enabling fluorescent detection of secretion across different secretion systems.

**Figure 6:**
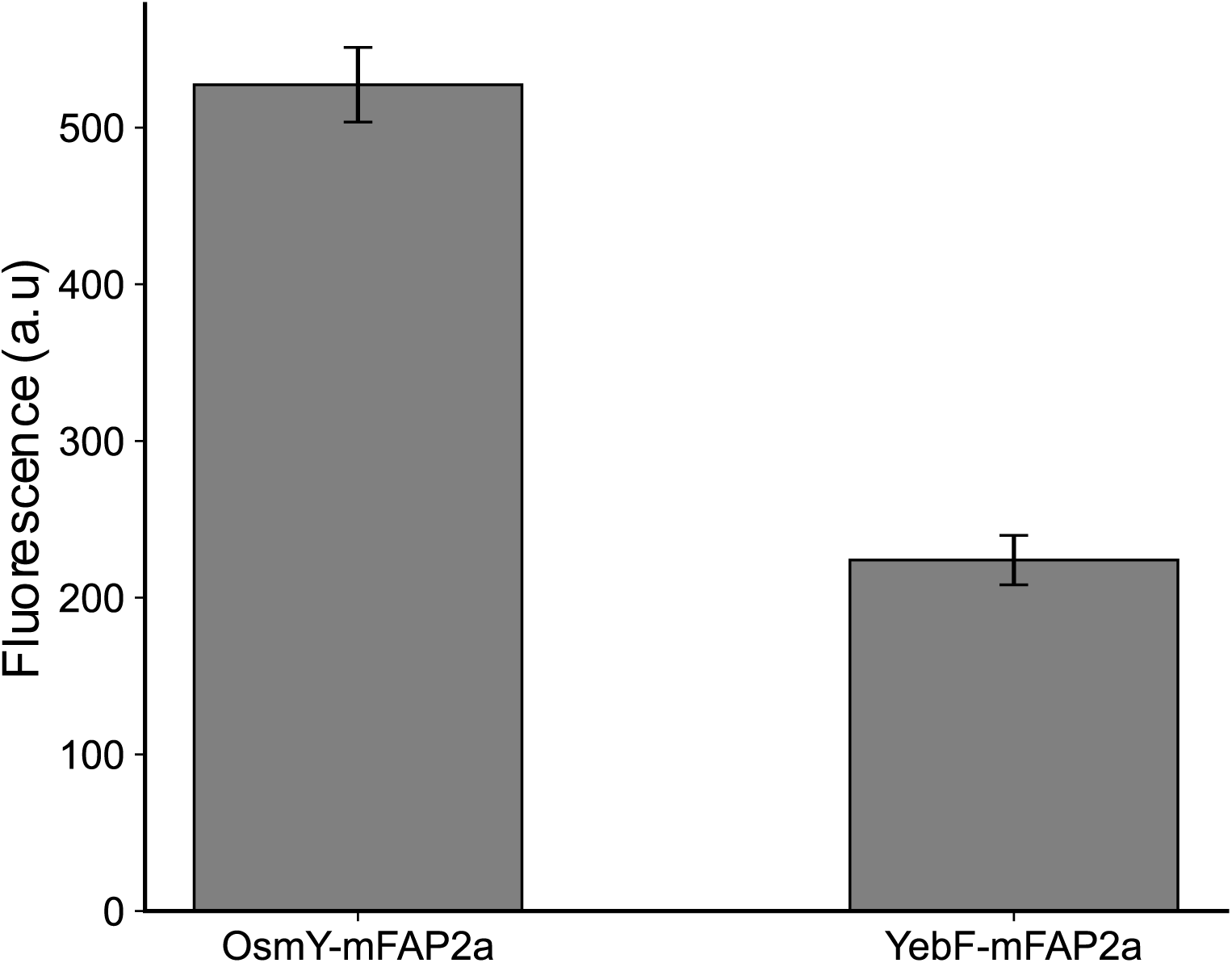
mFAP can be used to track secretion through other secretory pathways. *E. coli* strains BL21(DE3) harboring an expression plasmid for mFAP fused to either OsmY or YebF were grown and secretion samples were collected at the end of culture. Secretion samples were measured for background subtracted fluorescence using the mFAP assay. Error bars represent the standard error of the mean of three biological replicates.

## Discussion

This work demonstrates the power of mFAP as a fast, quantitative secretion assay. Its fluorescent signal develops quickly, remains stable, and is compatible with rich media, overcoming challenges faced by other fluorescent reporters. We showed its utility in measuring secretion through the T3SS of *S*. Typhimurium and fusion with *E. coli* proteins YebF and OsmY, suggesting it could serve as a universal reporter across secretion systems, potentially including eukaryotic systems.

We applied the mFAP assay for two purposes: assessing combinatorial strain designs for enhanced secretion and studying secretion dynamics under regulatory changes. This highlights just a fraction of the assay’s potential. For example, researchers could use mFAP to develop generalizable secretion tags that yield high titers across target systems (Gao, Wang, Li, Yu, & Qi, 2015; Han et al., 2017; Ling et al., 2017). The assay enables screening of large mutant libraries, which have potential to reveal functional insights such as those identified by mutating the PrgI protein (Burdette et al., 2024). All such studies require the generation of libraries with hundreds to thousands of mutants, which the mFAP assay is well-equipped to screen. However, while the assay can reasonably handle up to thousands of samples, future work incorporating whole genome approaches can contain tens of thousands to millions of library members, and being able to directly measure secretion titer with such a large sample set will require additional novel creative and innovative solutions.

## Materials and Methods

### Strains and Growth Conditions

All *S*. Typhimurium strains are derived from the strain ASTE13, which is derivative *S*. Typhimurium strain LT2 (Song et al., 2017a). All *E. coli* strains are derived from the BL21(DE3) strain. For expression and secretion experiments, all strains were grown from single colonies from fresh streaks of frozen glycerol stocks. Strains were grown at 37°C with shaking at 225 rpm in an orbital shaker overnight for 12-16 hours in the lysogeny broth Lennox formulation (LB-L, Dot Scientific, MI) for *S*. Typhimurium or lysogeny broth Miller formulation (LB-M, Fisher Scientific, NH) for *E. coli*. *S.* Typhimurium strains were subcultured 1:100 in LB-L, LB-L supplemented with 89 mM potassium phosphate and 0.4% glycerol (Burdette et al., 2021), or No Carbon Essential media (NCE; 1mM MgSO_4,_ 34 mM K_2_HPO_4_, 29 mM KH_2_PO_4_, 17 mM Na(NH_4_)HPO_4_, 0.05 mM ferric citrate, 0.4% glycerol) and induced with 100 µM isopropyl β-d-1-thiogalactopyranoside (IPTG) at the time of subculture unless no activation plasmid was present. Chloramphenicol (34 μg/mL) and/or kanamycin sulfate (50 μg/mL) were added to cultures as appropriate. Subculture volumes were either 5 mL in a 24 well block (Axygen, NY) or 0.5 mL in a 96 well block (Axygen, NY). *E. coli* strains were subcultured 1:100 in LB-M with appropriate antibiotics and induced with 100 µM IPTG when the OD reached ∼0.5. The strains were then grown for 8 hours at 37°C and 225 rpm in an orbital shaker. Samples of the whole culture were taken after 8 hours. Supernatant samples were taken from the culture after centrifugation at 4,500 x g for 10 minutes, which we refer to as the secreted fraction.

### DNA Manipulation

PCR was performed with the *Phusion* or *Vent* polymerases (New England Biolabs, MA). Constructs carrying the secretion chaperone, secretion tag, and mFAP gene (mFAP2a or mFAP10), 2x-FLAG tag, and 6x-His tag were assembled using the Golden Gate method (Engler, Gruetzner, Kandzia, & Marillonnet, 2009) using a previously developed backbone containing the native P_sic_ promoter (Widmaier et al., 2009a). pET28b-YebF-mFAP2a and pET28b-OsmY-mFAP2a were also assembled using the Golden Gate method. All cloning was performed in the *E. coli* strain DH10B. All DNA sequences were verified by Sanger sequencing (Quintara, CA). A table of strains used in this study are provided in Table 1.1 in the supplementary information. A list of plasmids and primers used in this study are provided in Table 1.2 and 1.3, respectively, in the supplementary information. Sequence-verified plasmids were used to transform electrocompetent strains. Successful transformants were identified following selection on LB-antibiotic plates.

### Protein separation and western blotting

Whole culture samples were diluted 1:2 in 4x Laemmli buffer with 10% β-mercaptoethanol and boiled at 95°C for 5 minutes. Supernatant samples were diluted 1:6 in 1x Laemmli with 10% β-mercaptoethanol and boiled at 95°C for 5 minutes, except for SptP^167^-mFAP2a in enhanced LB-L, which was diluted 1:24 in 1x Laemmli. Samples were separated by SDS-PAGE and then transferred to a nitrocellulose membrane (Millipore, MO) using the TransBlot SD unit (Bio-Rad, CA). Membranes were interrogated with a mouse monoclonal anti-FLAG antibody (Sigma-Aldrich, MO) (1:6666) and then a secondary anti-Mouse-HRP-IgG (Fisher Scientific, MA) (1:1000) antibody. Bands were visualized using the Pierce ECL substrate (Fisher Scientific, MA) and imaged with a Chemidoc XRS+ unit (Bio-Rad, CA) or a Workhouse Gel Imager (Azure Biosystems, CA). Relative protein amounts were corrected for dilution if appropriate. Relative protein quantities were determined using densitometry analysis in Image Lab software (Bio-Rad, CA). Values were normalized to the average of the replicates for the designated normalization condition.

### mFAP fluorescence assay

100 μL whole culture samples were combined with 100 μL of 200 µM DFHBI for at least 10 minutes. 100 μL supernatant samples or pure protein samples diluted in the supernatant of cultures harboring no secretion plasmid were combined with 100 μL of 100 µM DFHBI for 10 minutes unless otherwise noted. Fluorescence of samples was measured at 467 nm absorbing and 505 nm emitting using a Synergy H1 plate reader with monochrometer (Biotek, VT) and Gen5 software (Biotek, VT) with a gain of 75.

### Protein purification from *Salmonella* supernatant

SptP^167^-mFAP2a-2xFLAG-6xHis was expressed from a plasmid under the control of the native T3SS promoter P_sic_ and purified from the ASTE13 strain of *S.* Typhimurium. Briefly, a single colony from a fresh streak of a frozen glycerol stock was grown in LB-L overnight at 37°C and 225 rpm in an orbital shaker. The overnight culture was then subcultured 1:100 into 50 mL Terrific Broth (Fisher Scientific, MA) and grown for 8 hours at 37°C and 225 rpm in an orbital shaker. Cells were harvested by centrifugation at 4,500 x g for 10 minutes. The supernatant was passed through a His Gravitrap Column (GE Healthcare, IL) and washed three times with 10 mL binding buffer (20 mM NaPO_4_, 0.5 M NaCl, 20 mM imidazole, pH 8). The protein was then eluted in 3.5 mL elution buffer (20 mM NaPO_4_, 0.5 M NaCl, 500 mM imidazole, pH 8) and buffer exchanged into Tris-buffered saline (25 mM Tris, 100 mM NaCl, pH 8) using a PD10 column (GE Healthcare, IL). The protein was quantified using a Nanodrop (Thermo Scientific, MA), and the purity was measured via SDS-PAGE and Coomassie staining.

### Recombineering

(Adapted from Burdette et al., 2021) Recombineering was performed in *S*. Typhimurium ASTE13 as described by Thomason et al. (Thomason et al., 2007). In summary, a *cat-sacB* cassette conferring chloramphenicol resistance and sucrose sensitivity was amplified from the *E. coli* TUC01 genome using primers with 40 bp of homology 5′ and 3′ to the locus of interest (*invE, sipBCD, hilE,* and *prgI*). The *prgI* gene with the S49R mutation was amplified using primers containing the same 40 bp of homology 5′ and 3′ to the locus of interest as used for the *cat-sacB* cassette. PCR was performed with Phusion DNA polymerase. *S.* Typhimurium ASTE13 was first transformed with pSIM6. The first round of recombineering inserted the *cat-sacB* cassette at the locus of interest, and the second round replaced the *cat-sacB* cassette with the target DNA product. The replacement DNA was either the *prgI::S49R* PCR product for the *prgI* locus or a 200 bp double-stranded DNA insert containing the first and last 30 bp of the gene to be deleted (either *invE, sipBCD,* or *hilE*) flanked 5′ and 3′ by 70 bp of homology to the gene’s locus. Sanger sequencing (Quintara, CA) confirmed the genomic modifications, and the strains were subsequently cured of pSIM6.

## Abbreviations

T3SS: Type 3 Secretion System
mFAP: mini-Fluorescent Activating Protein
DFHBI: 5Z)-5-[(3,5-Difluoro-4-hydroxyphenyl)methylene]-3,5-dihydro-2,3-dimethyl-4H-imidazol-4-one
LB-L: Lysogeny broth, Lennox formulation
LB-M: Lysogeny broth, Miller formulation
LB-ES: Lysogeny Broth (Lennox formulation), enhanced secretion
NCE: No Carbon Essential medium
IPTG: isopropyl β-d-1-thiogalactopyranoside

## Supplementary material description

Supplemental figures/tables consist of a variety of fluorescence data, standard curves, and western blots intended to strengthen the observations made in the main results section.

## Acknowledgements

We want to acknowledge and thank Dr. David Baker and Dr. Jason Klima for providing us with input and expertise on the mFAP protein. We would like to thank colleagues in the research group of Dr. Michael Jewett for allowing us to use their plate reader. SAL and DTE were supported by the National Science Foundation (award number BBE-1706125). JSS and DTE were supported by National Science Foundation (award number DMR-2308691) and the National Institutes of Health Training Grant (T32GM008449) through Northwestern University’s Biotechnology Training Program.

## Author’s Contributions (CRediT)

**Samuel A. Leach:** Conceptualization (lead); data curation (lead); formal analysis (lead); funding acquisition (equal); investigation (lead); methodology (equal); writing original draft preparation (lead). **Jordan Summers:** Conceptualization (supporting); data curation (supporting); formal analysis (supporting); funding acquisition (equal); investigation (supporting); methodology (equal); software (lead); validation (lead); visualization (lead); writing original draft & preparation (supporting); writing review & editing (lead). **Edward Wen:** Resources (supporting). **Danielle Tullman-Ercek:** Conceptualization (supporting); formal analysis (supporting); funding acquisition (equal); methodology (supporting); project administration (lead); resources (lead); supervision (lead); writing – review & editing (equal).

