## Supplementary Figures for "A High Throughput Assay for Measuring Secreted Protein Based on a *de novo* Fluorescent Reporter Reveals Regulatory and Structural Insights in *Salmonella* Type Three Secretion System"

**Supplementary Material**

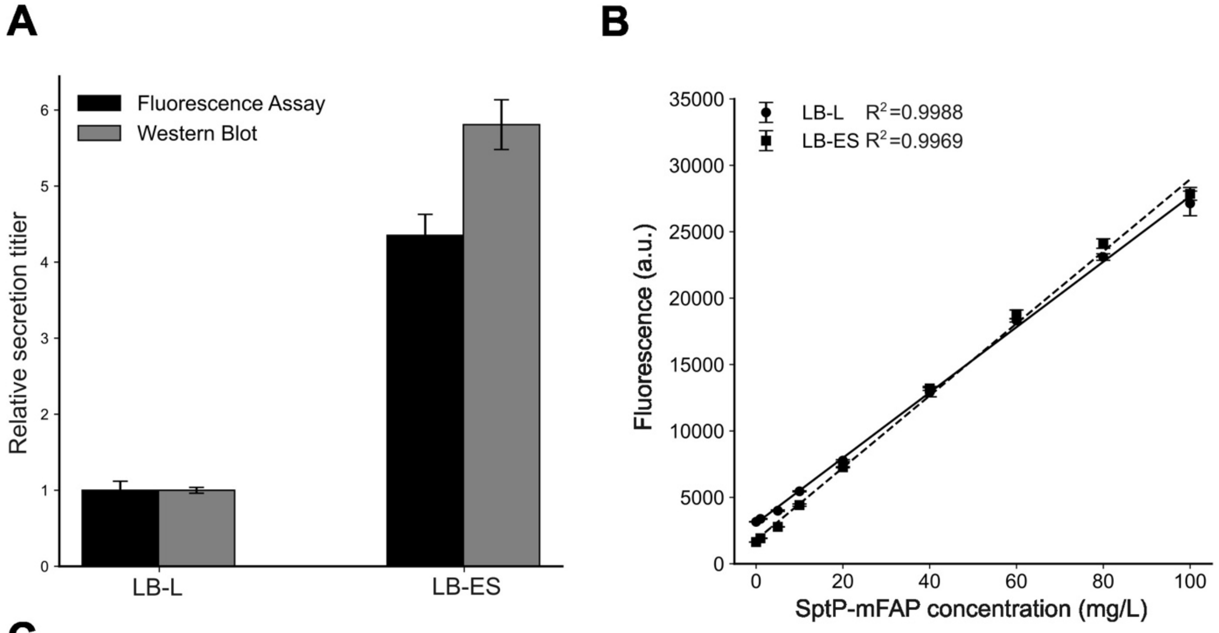


C

| SptP-mFAP concentration (mg/L) | % fluorescence variation over time | % fluorescence variation between t=10 min and t=90 min |
|---|---|---|
| 10 | 1.37 | 10.16 |
| 20 | 0.91 | 3.79 |
| 40 | 2.53 | 3.66 |
| 60 | 1.10 | 7.25 |
| 80 | 1.13 | 9.34 |
| 100 | 4.71 | 13.19 |

**Supplementary Figure 1:** mFAP concentration and fluorescence agree over various conditions. A) Relative secretion titer from the mFAP assay and western blotting in both LB and LB-ES. B) mFAP fluorescence remains consistent across a wide concentration for both LB and LB-ES. C) Average percentage fluorescence variation over time and between t=10 minutes and t=90 minutes for varying concentrations of SptP-mFAP.

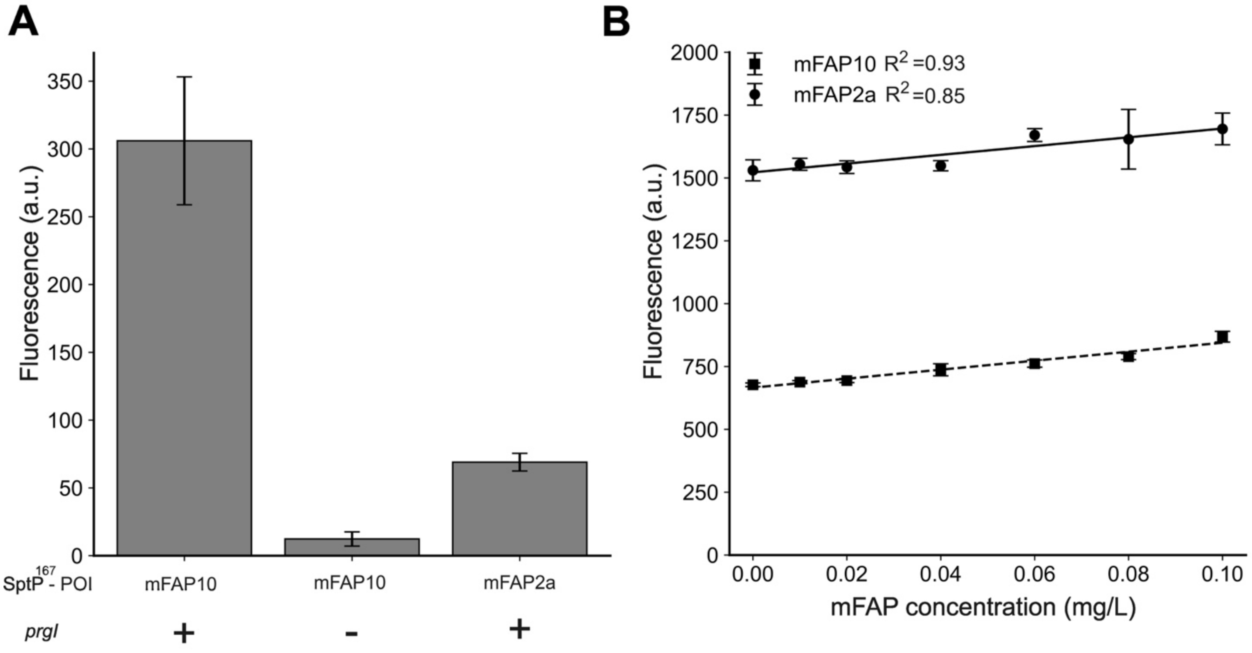


**Supplementary Figure 2:** Comparison of mFAP2a and mFAP10 fluorescence. A) mFAP assay of mFAP10 with and without *prgI* and mFAP2a fused to SptP$^{167}$. B) mFAP10 displays more robust linearity at >1 mg/L concentrations compared to mFAP2a.

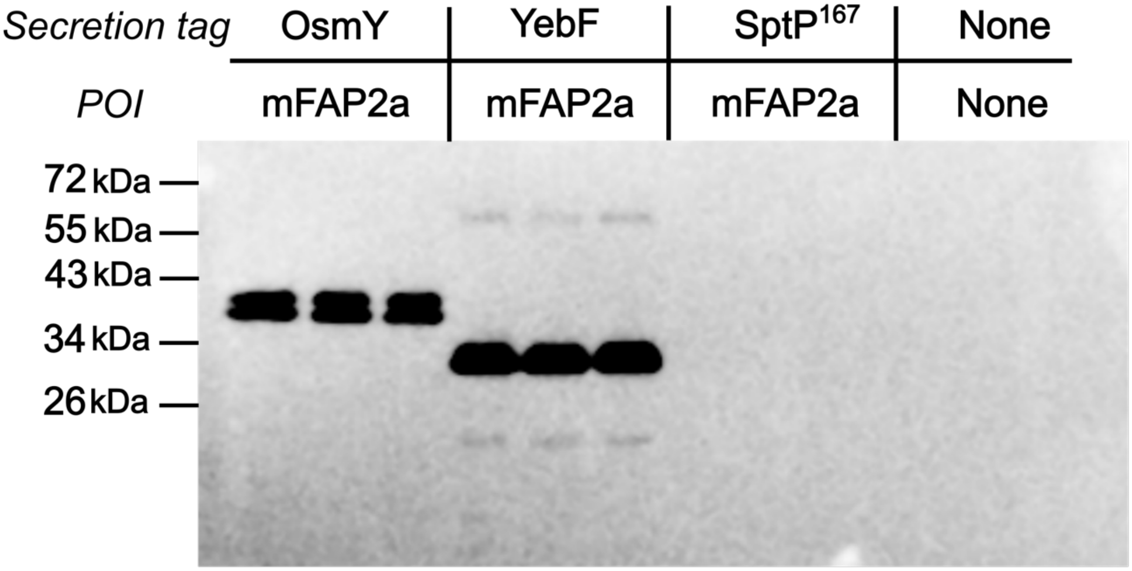


**Supplementary Figure 3:** mFAP is selectively secreted by OsmY and YebF. Secreted fractions of BL21 (DE3) harboring plasmids encoding for either OsmY, YebF, or SptP N-terminal fusions to mFAP2a-2xFLAG were collected and blotted for the FLAG epitope. SptP-mFAP2a is not detected in the secreted fraction, while empty BL21 (no secretion tag or POI) suggests no non-specific interactions.
