## Supplemental Tables for "A High Throughput Assay for Measuring Secreted Protein Based on a *de novo* Fluorescent Reporter Reveals Regulatory and Structural Insights in *Salmonella* Type Three Secretion System"

### Table 1.1 Strains Used in This Work

| Strain Name | Organism | Comment | Reference |
|---|---|---|---|
| ASTE13 | S. Typhyimurium | LT2-derived lab strain similar to DW01 | (Song et al., 2017) |
| ASTE13 Δ*prgI* | S. Typhyimurium | lacks T3SS needle monomer; secretion incompetent | (Metcalf, Finnerty, Azam, Valdivia, & Tullman-Ercek, 2014) |
| ASTE13 Δ*hilE* | S. Typhyimurium | Lacks T3SS repressor HilE | (Liang, Burdette, Wong, & Tullman-Ercek, 2023) |
| BL21(DE3) | *E. coli* | | (Jeong, Kim, & Lee, 2015) |
| ASTE13 *prgI::S49R* | | mutation in T3SS needle monomer conferring enhanced secretion | (Burdette et al., 2024) |
| ASTE13 Δ*sipBCD* | | lacks T3SS tip complex | This study |
| ASTE13 Δ*invE* | | lacks T3SS structural regulator | This study, based on(Kubori & Galán, 2002) |
| ASTE13 *prgI::S49R,* Δ*sipBCD* | | | This study |
| ASTE13 *prgI::S49R,* Δ*invE* | | | This study |
| ASTE13 *prgI::S49R,* Δ*sipBCDinvE* | | | This study |
| ASTE13 *prgI::S49R,* Δ*sipBCDinvEhilE* | | | This study |
| ASTE13 *prgI::S49R,* Δ*sipBCDhilE* | | | This study |

### Table 1.2 Plasmids Used in This Work

| Plasmid Name | ORFs Under Control | ORI | abR | Reference |
|---|---|---|---|---|
| Psic SptP-mFAP | *sicP; sptP-mFAP2a-2XFLAG-6xHis* | colE1 | cam | This study; (Widmaier et al., 2009) |
| PlacUV5 HilA | *hilA* | p15a | kan | (Metcalf et al., 2014) |
| Psic SptP-DH | *sicP; sptP-DH-2XFLAG-6xHis* | colE1 | cam | (Widmaier et al., 2009) |
| Psic SopB-mFAP | *sigE; sopB-mFAP2a-2XFLAG-6xHis* | colE1 | cam | This study; (Widmaier et al., 2009) |
| Psic SipC-mFAP | *sicA; sipC-mFAP2a-2XFLAG-6xHis* | colE1 | cam | This study; (Widmaier et al., 2009) |
| Psic SipA-mFAP | *invB; sipA-mFAP2a-2XFLAG-6xHis* | colE1 | cam | This study; (Widmaier et al., 2009) |

| | | | | |
|---|---|---|---|---|
| Psic SopA-mFAP | *invB; sopA-mFAP2a-2XFLAG-6xHis* | colE1 | cam | This study; (Widmaier et al., 2009) |
| Psic SopE2-mFAP | *invB; sopE2-mFAP2a-2XFLAG-6xHis* | colE1 | cam | This study; (Widmaier et al., 2009) |
| Psic SopD-mFAP | *sopD-mFAP2a-2XFLAG-6xHis* | colE1 | cam | This study; (Widmaier et al., 2009) |
| Psic SptP-mFAP10 | *sicP; sptP-mFAP10-2XFLAG-6xHis* | colE1 | cam | This study; (Widmaier et al., 2009) |
| pET28b YebF-mFAP | *yebF-mFAP2a* | colE1 | kan | This study; |
| pET28b OsmY-mFAP | *osmY-mFAP2a* | colE1 | kan | This study, based on(Gonzalez-Perez et al., 2021) |

### Table 1.3 Primers Used in This Work

| Primer Sequence | Amplicon | Used to Construct |
|---|---|---|
| TAGGGAGGTCTCGGCTTTCACGCGCCGCGCAGCTC | *mFAP2a; mFAP10* | Psic SptP-mFAP; Psic SptP-mFAP10 |
| CTGAGTGGTCTCGCGCTTTCCTGACGCTGCAGGGTACC | *mFAP2a; mFAP10* | Psic SptP-mFAP; Psic SptP-mFAP10 |
| TGTCACGGTCTCGATTCGAGTCTTGAGGTAACTATATGGAAAGTCTATTAAAT | *sigE sopB* | Psic SopB-mFAP |
| TTTAATCGCGCTCCTGATGGCGAACTGGGGATATTATGCTTAATATTCAATGTGACGGAAGATCACTTCG | *cat-sacB* | ASTE13 Δ*sipD* (Burdette, Wong, & Tullman-Ercek, 2021) |
| CTTACACTTGTAACCATTATTAATATCCTCTTCTGTTATCCTTGCAGGAAATCAAAGGGAAAACTGTCCATAT | *cat-sacB* | ASTE13 Δ*sipD* (Burdette et al., 2021) |
| TCTGAAAGGTCATCTATACGCCATCATGGGTGTGATTTAATCGCGCTCCTGATGGCGAACTGGGGATATTATGCTTAATATTCAATTCCTGCAAGGATAA | *sipD* | ASTE13 Δ*sipD* (Burdette et al., 2021) |
| TCTGCATACCTGGCATTATGACGGGGGGCTGAGTCCTTACACTTGTAACCATTATTAATATCCTCTTCTGTTATCCTTGCAGGAATTGAATATTAAGCAT | *sipD* | ASTE13 Δ*sipD* (Burdette et al., 2021) |
| CGCTAAAAACGGCGGAGACAGAGCAGCACAGTGAACAAGAAAAGGAATAATGTGACGGAAGATCACTTCG | *cat-sacB* | ASTE13 Δ*sipBCD* |

| | | |
|---|---|---|
| ACGGGGGGCTGAGTCCTTACACTTGTAA CCATTATTAATATCCTCTTCTGATCAAAG GGAAAACTGTCCATATG | *cat-sacB* | ASTE13 Δ*sipBCD* |
| TGGTCTATCTGGAGGCGCTAAAAACGGCG GAGACAGAGCAGCACAGTGAACAAGAAA AGGAATAACAGAAGAGGATATTAATAAT GG | *sipBCD* | ASTE13 Δ*sipBCD* |
| GGTCTGCATACCTGGCATTATGACGGGGG GCTGAGTCCTTACACTTGTAACCATTATTA ATATCCTCTTCTGTTATTCCTTTTCTTG | *sipBCD* | ASTE13 Δ*sipBCD* |
| TAATATTAAATTAAGACAGCTTTTCAATAG TACGACGCTGTGTGACGGAAGATCACTTCG | *cat-sacB* | ASTE13 Δ*invE* |
| AGGCAATTAAATGATTCCTGGCTCAACC TCCGGTATTTCAATCAAAGGGAAAACTG TCCATAT | *cat-sacB* | ASTE13 Δ*invE* |
| TTAAATTAAGACAGCTTTTCAATAGTACG ACGCTGTGAAATACCGGAGGTTGAGCCA GGAATCATTTAAT | *invE* | ASTE13 Δ*invE* |
| ACGAAATGGCTGGAAAATGGAACGTTCT TTCATTGTTGGCTGTGACGGAAGATCACT TCG | *cat-sacB* | ASTE13 Δ*hilE* |
| GTCCTCATCGCCACAGCGCCTGTCGGTGA AGAGGCCGCCATCAAAGGGAAAACTGTC CAT | *cat-sacB* | ASTE13 Δ*hilE* |
| ATGGCTGGAAAATGGAACGTTCTTTCATT GTTGGCGGCGGCCTCTTCACCGACAGGCG CTGTGGCGATGA | *hilE* | ASTE13 Δ*hilE* |
| TGTCACGGTCTCGCGCTGTTATTAAGCT GCTTGACCTG | *sigE sopB* | Psic SopB-mFAP |
| TGTCACGGTCTCGATTCACAGATAACAG GAGTAAGTAATGGATTATCAAAA | *sicA sipC* | Psic SipC-mFAP |
| TGTCACGGTCTCGCGCTTCCGCTAATATC AAAAAACTTTCCGACTTC | *sicA sipC* | Psic SipC-mFAP |
| TGTCACGGTCTCGATTCTTAATTAAGGAA AAGATCTATGCAACATTTG | *invB sipA; invB sopE2; invB sopA* | Psic SipA-mFAP; Psic SopE2 mFAP; Psic SopA-mFAP |
| TGTCACGGTCTCGCGCTACCGCCAGTGTT ATTTTTGATAATATCTAACAGC | *invB sipA* | Psic SipA mFAP |
| TGTCACGGTCTCGCGCTGGCCGGATCTTT ACTCGCATTA | *invB sopE2* | Psic SopE2-mFAP |
| TGTCACGGTCTCGTCCTTTATCTCATTAGC GACCG | *invB SopA* | Psic SopA-mFAP |

| | | |
|---|---|---|
| TGTCACGGTCTCGATTCATTTGAAGGAAAATATTATGCCAGTCAC | *sopD* | Psic SopD |
| TGTCACGGTCTCGCGCTGACTTTATCCCAACCCCCCCATA | *sopD* | Psic SopD |
| GTCTCAGGTCTCCCATGATGAAAAAAAGAGGGGCGTTTTTAGGG | *yebF* | pET28b YebF-mFAP |
| CTGCGCGGCGCGTGAACGCCGCTGATATTCCGC | *yebF* | pET28b YebF-mFAP |
| GCGGAATATCAGCGGCGTTCACGCGCCGCGCAG | *mFAP2a* | pET28b YebF-mFAP |
| GTCTCAGGTCTCCCTTAGTGGTGATGGTGATGATGCTTGTCA | *mFAP2a* | pET28b YebF-mFAP; pET28b OsmY-mFAP |
| TTTCCCGGTCTCGCATGATGACTATGACAAGACTGA | *osmY* | pET28b OsmY-mFAP |
| CTGCGCGGCGCGTGACTTAGTTTTCAGATCATTTTTAAC | *osmY* | pET28b OsmY-mFAP |
| GTTAAAAATGATCTGAAAACTAAGTCACGCGCCGCGCAG | *mFAP2a* | pET28b OsmY-mFAP |
